# R-spondin1 regulates muscle progenitor cell fusion through control of antagonist Wnt signaling pathways

**DOI:** 10.1101/063669

**Authors:** Floriane Lacour, Elsa Vezin, Florian Bentzinger, Marie-Claude Sincennes, Robert D. Mitchell, Ketan Patel, Michael A. Rudnicki, Marie-Christine Chaboissier, Anne-Amandine Chassot, Fabien Le Grand

## Abstract

Tissue regeneration requires the selective activation and repression of specific signaling pathways in stem cells. As such, the Wnt signaling pathways have been shown to control stem cell fate. In many cell types, the R-Spondin (Rspo) family of secreted proteins acts as potent activators of the canonical Wnt/β-catenin pathway. Here, we identify Rspo1 as a mediator of skeletal muscle tissue repair. Firstly we show that *Rspo1-null* muscles do not display any abnormalities at the basal level. However deletion of *Rspo1* results in global alteration of muscle regeneration kinetics following acute injury. We found that muscle stem cells lacking *Rspo1* show delayed differentiation. Transcriptome analysis further demonstrated that Rspo1 is required for the activation of Wnt/β-catenin target genes in muscle cells. Furthermore, muscle cells lacking *Rspo1* fuse with a higher frequency than normal cells, leading to larger myotubes containing more nuclei both *in vitro* and *in vivo*. We found the increase in muscle fusion was dependent on up-regulation of non-canonical Wnt7a/Fzd7/Rac1 signaling. We conclude that antagonistic control of canonical and non-canonical Wnt signaling pathways by Rspo1 in muscle stem cell progeny is important for restitution of normal muscle architecture during skeletal muscle regeneration.

## INTRODUCTION

Adult muscle stem cells, called satellite cells (MuSCs), located around the differentiated myofibers exist in a quiescent state and are readily identified through their expression of the Paired-Box transcription factor Pax7 (Seale et al., 2000). Following injury to the host myofibers they become activated and proliferate give rise to muscle progenitor cells expressing the myogenic regulatory factors MyoD and Myogenin which act to promote differentiation (Chargé and Rudnicki, 2004). Differentiating myocytes will then fuse together to form myotubes that will further mature into new myofibers. Myocyte differentiation and fusion processes are highly regulated (Abmayr and Pavlath, 2012). Perturbations in migration, adhesion or membrane fusion can give rise to defects in the reconstitution of muscle architecture. As such, accelerated fusion can lead to the formation of giant dystrophic myofibers (Charrin et al., 2013) while a reduction in myocyte fusion potential results in smaller muscles with myofibers containing fewer nuclei (Horsley et al., 2003). This is exemplified in skeletal muscle pathologies such as Duchenne Muscle Dystrophy (DMD). In dystrophic muscles, the continuous regeneration process led to the formation of abnormal fibers, termed branched or split fibers, which are more prone to damage during contraction (Chan et al., 2007), lack force generation, and contribute to the disease progression (Head, 2010).

The WNT signaling pathways are crucial regulators of adult muscle regeneration (von Maltzahn et al., 2012). The canonical Wnt/β-catenin pathway is required for muscle progenitor cell differentiation and regulate both skeletal muscle development (Borello et al., 2006, Anakwe et al., 2003) and repair following injury (Brack et al., 2009, Parisi et al., 2015). In contrast, the non-canonical Wnt7a/Fzd7 signaling pathway stimulates skeletal muscle growth and repair by inducing the symmetric expansion of satellite stem cells through the planar cell polarity (PCP) pathway (Le Grand et al., 2009) and by activating the anabolic Akt growth pathway in myofibers (von Maltzahn et al., 2012). Recently it was shown that Wnt7a/Fzd7/Rac1 signaling enhance MuSCs and myoblasts migration (Bentzinger et al., 2014). In many tissues, both canonical (Carmon et al., 2011) and non-canonical (Glinka et al., 2011) Wnt pathways can be overstimulated by R-spondins. The R-spondin family is comprised of four different secreted proteins (RSPO1-4) (de Lau et al., 2012). Each R-spondin has its own expression pattern and properties. R-spondins interact with leucine-rich repeat-containing G protein-coupled *receptors* (*LGR*) (de Lau et al., 2011). LGRs allow a potentiation of the Wnt signaling pathways in response to R-spondins by neutralizing the E3 ligases RNF43 and ZNRF3 which act to remove Wnt receptors from the stem cell surface (Koo et al., 2012, Hao et al., 2012).

Number of studies reported the essential roles played by R-spondins both during embryonic development, adult tissue homeostasis and repair. Thus, *Rspo1* is required for female sexual development and *Rspo1-null* mice show masculinized gonads (Parma et al., 2006, Chassot et al., 2008). In adult tissues, RSPO1 promotes growth of the intestinal epithelium (Kim et al., 2005) and is a beta-cell growth factor in the pancreas (Wong et al., 2010). Moreover, Lgr5/Rspo1-expressing stem cells are able to regenerate hepatocytes and bile ducts upon liver damage *in vivo* (Huch et al., 2013). To date, the role of R-spondins in regenerative myogenesis remains unclear. It is known that R-spondins are also expressed in muscle cells and the C2C12 myoblast cell line has been shown to express *Rspo2* and *Rspo3.* In these cells, RSPO2/LGR4 regulate myogenic differentiation (Han et al., 2011). A recent report documented that *Rspo1* transcript expression was strongly up-regulated in primary myoblasts over-expressing PAX7 compared to control cells (Soleimani et al., 2012). However, so far the role of R-spondins in regenerative myogenesis remains undetermined. Here we aimed to investigate the function of *Rspo1* in steady-state muscle as well as during regeneration.

## RESULTS

### Muscle satellite cells express the Pax7 target gene *Rspo1*

To further validate the expression of Rspo1 in adult MuSCs, we performed immunocytochemical analysis for PAX7 and RSPO1 on myofibers isolated from *Extensor Digitorum Longus* (EDL) muscles. We observed that quiescent MuSCs expressed RSPO1 proteins and that activated MuSCs on cultured myofibers (24 hours after isolation) expressed relatively higher levels of RSPO1 proteins (Figure 1A). Of note, RSPO1 is mostly cytoplasmic in quiescent MuSCs and appears both cytoplasmic and nuclear in activated MuSCs. Analysis of primary myoblasts cultured *in vitro* further confirmed the expression of RSPO1 by differentiating muscle progenitor cells (Figure 1B).

**Figure 1.**
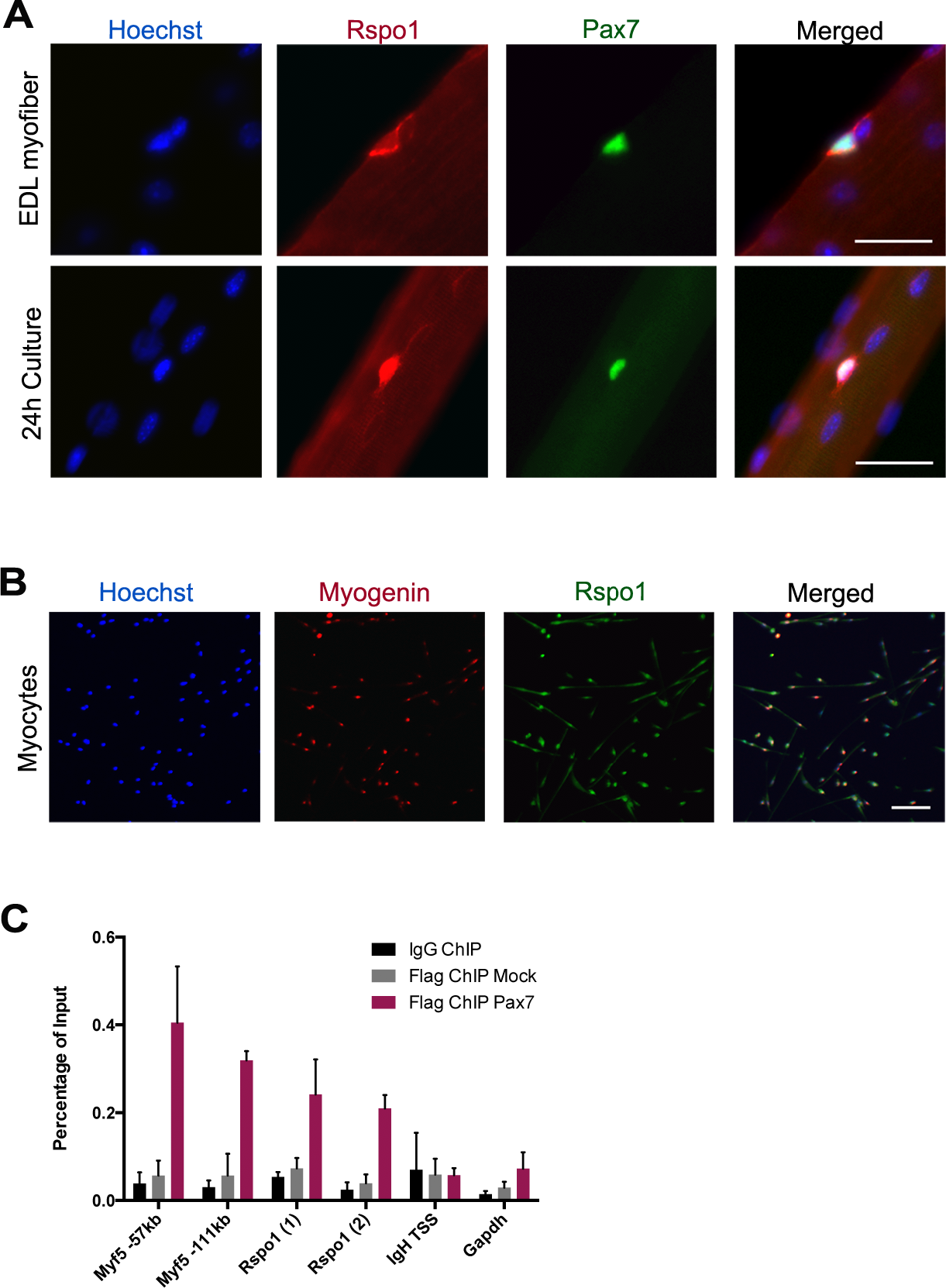
R-spondin1 is expressed in MuSCs and primary myocytes. (A) Immunolocalization of RSPO1 (red) and PAX7 (green) proteins in MuSCs on single myofibers. Nuclei are stained with Hoechst (blue). (B) Immunolocalization of MYOGENIN (red) and RSPO1 (green) proteins in 24h-differentiated myocytes. Nuclei are stained with Hoechst (blue). (C) Occupancy of PAX7-Flag proteinx at the promoters of *Myf5, Rspo1* and *Gapdh* (control) genes in primary myoblasts. The input represents the relative enrichment of PAX7-Flag proteins compared to the controls (IgG and Flag-Mock). Bars: 20 μm

Since *Rspo1* expression was shown to be strongly up-regulated in myoblasts expressing Pax7-Flag, we tested if *Rspo1* is a direct target of *Pax7* in adult myogenic cells. To this aim, we asked whether PAX7 proteins can bind to a putative binding site localized 35kb upstream of *Rspo1* gene identified by Chromatin Immunoprecipitation (ChIP) coupled with deep sequencing (Soleimani et al., 2012). ChIP assays were thus performed using primary myogenic cell cultures expressing Pax7-Flag. They demonstrated that PAX7 proteins were bound close to the *Rspo1* gene as well as to the positive controls (*Myf5 -111kb and -57kb* enhancer regions), as relative to a mock IP and to the negative controls (IgH and Gapdh loci) (Figure 1C). These results indicate that *Rspo1* is a direct target of PAX7 in adult muscle progenitor cells.

### *Rspo1* is not required for muscle tissue formation

We therefore investigated whether Rspo1 plays a role in the adult skeletal muscle tissue. To this aim, we bred constitutive *Rspo1* knock-out mice (Chassot et al., 2008). Homozygous mutant mice present a complete *Rspo1* loss-of-function leading to female-to-male sex reversal and sterility in females, otherwise heterozygous mice are phenotypically normal and fertile. We did not observe any phenotypic difference in skeletal muscle histology of young adult (8 weeks-old) *Rspo1-null* compared to control mice (controls are wild-type and *Rspo1-heterozygous* mice) as shown by hematoxylin and eosin staining of *Tibialis Anterior* (TA) muscles (Figure S1A) and by measuring the weights of multiple limb muscles (Figure S1B). In TA muscles, loss of *Rspo1* expression did not influence the numbers of myofibers (Figure S1C), their size (Figure S1D) and the crosssectional area (CSA) distribution (Figure S1E). In agreement with these findings, quantification of the number of myonuclei per myofiber (Figure S1F) and of the entire muscle CSA (Figure S1G) did not show any significant difference between the two genotypes. Furthermore, *Rspo1-null* and control animals showed similar number of Pax7-expressing MuSCs on TA (Figure S1H) Soleus, Gastrocnemius and Triceps (data not shown) cryosections. Our analysis suggests that *Rspo1* is not required for myofiber formation and cellular organization or for the establishment of the MuSC population.

### *Rspo1* regulates skeletal muscle tissue regeneration

To address whether *Rspo1* is involved in skeletal muscle regeneration, we injured TA muscles from control and *Rspo1-null* mice by cardiotoxin injection (Vignaud et al., 2007). During the early phase of tissue regeneration (4d.p.i.) (Figure 2A), the number of Pax7+ progenitors was similar between control and *Rspo1-null* muscles (Figure 2B), but *Rspo1-null* muscles contained decreased numbers of Myogenin+ differentiating cells (Figure 2C) and of nuclei incorporated in myotubes compared to control muscles (Figure 2D), suggesting a delay in the differentiation process in *Rspo1-null* tissue. Later, at 7 d.p.i, (Figure 2E), newly formed myofibers were of similar caliber between both genotypes (Figure 2F) and the numbers of Myogenin+ cells inside myofibers was increased in *Rspo1-null* muscles compared to control muscles (Figure 2G), suggesting a compensatory enhancement of fusion in *Rspo1-null* muscles. At this time point, the number of Pax7+ progenitors remained similar between *Rspo1-null* and control muscles (Figure 2H).

**Figure 2.**
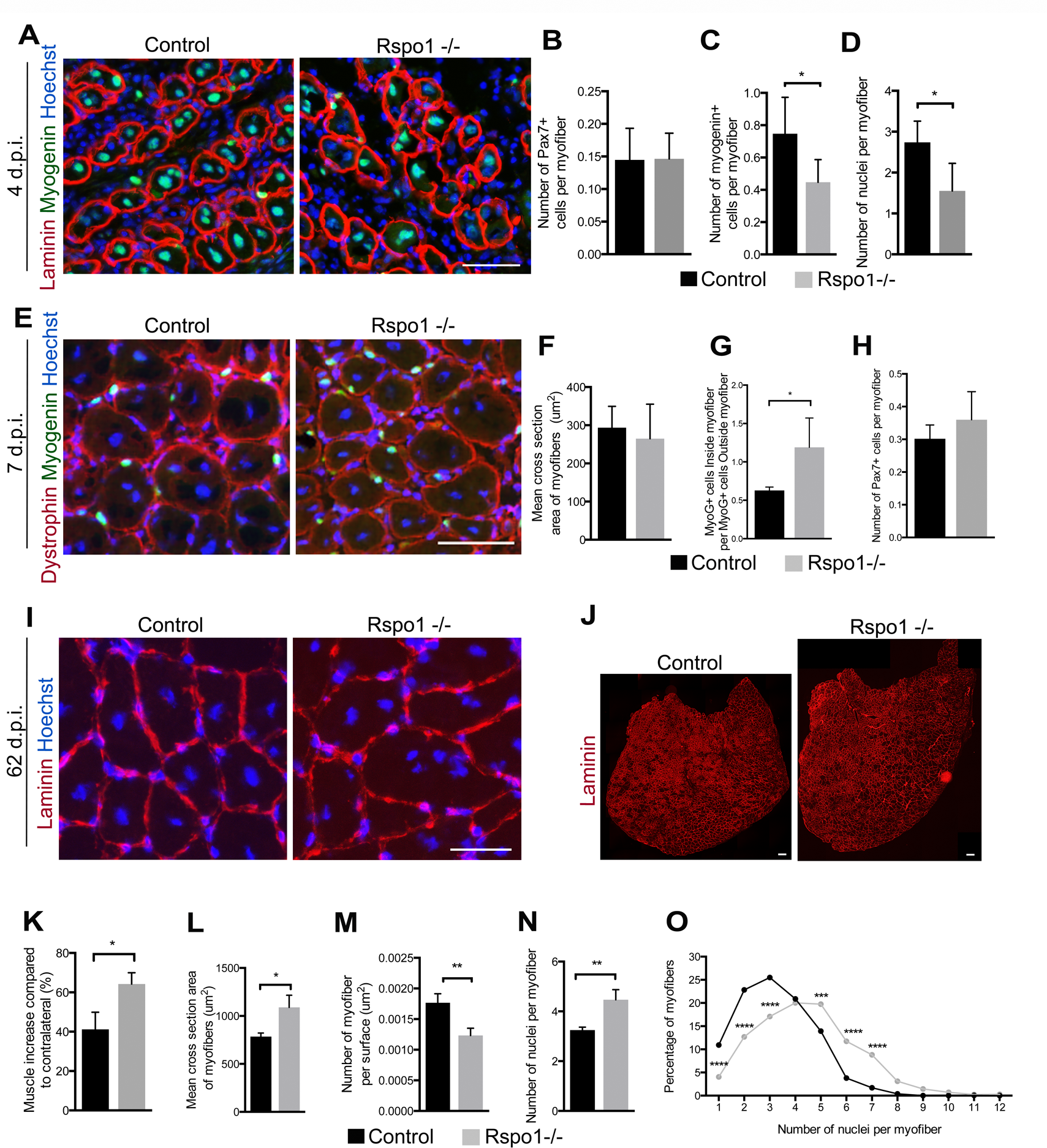
*Rspo1-null* muscles show enhanced regeneration caused by a delay of muscle progenitor cell differentiation and an improved fusion. (A) Immunolocalization of LAMININ (red) and MYOGENIN (green) proteins in control and *Rspo1-null* TA muscles at 4d.p.i. Nuclei are stained with Hoechst (blue). (B) Quantification of the number of PAX7-positives cells per myofiber 4 d.p.i. (C) Quantification of the number of cells expressing MYOGENIN per myofiber at 4 d.p.i. showing an alteration of the number of *Rspo1-null* differentiated cells. (D) Quantification of the number of nuclei per myofiber at 4 d.p.i. (E) Immunostaining of DYSTROPHIN (red) and MYOGENIN (green) proteins at 7 d.p.i. Nuclei are stained with Hoechst (blue). (F) Mean cross sectional area of myofibers in TA muscles showing no difference between control and *Rspo1-null* muscles at 7d.p.i. (G) Quantification of the number of MYOGENIN-positives cells inside the myofibers normalized by the number of cells outside the myofibers, showing an higher proportion of fused nuclei at 7 d.p.i. in *Rspo1-null* mice. (H) Quantification of the number of PAX7-positives cells per myofiber at 7 d.p.i. (I) Immunostaining of LAMININ (red) proteins on 62 d.p.i muscle sections. Nuclei are stained with Hoechst (blue). (J) Whole cross-section of 62 d.p.i. muscles. (K) Quantification of the TA muscle weight at 62 d.p.i. (L) Mean cross sectional area of the TA muscles at 62d.p.i. (M) Number of myofibers per surface at 62 d.p.i. (N) Quantification of the number of nuclei per myofiber at 62 d.p.i. (O) Distribution of the percentage of myofibers depending on their nuclei number at 62 d.p.i. Bars: 50 μm in A, E; 35 μm in I; 150 μm in J. *Error bars indicate standard deviation. *: p value <0.05.* **: *p value <0.01*

To further evaluate the effect of *Rspo1* mutation on muscle repair, we analyzed the morphology and cellular composition of regenerated muscles, 62 days after injury (Figure 2I). Strikingly, regenerated *Rspo1-null* muscles were larger (Figures 3J) and heavier (Figure 3K) compared to control muscles. Regenerated *Rspo1-null* muscles were composed of larger myofibers (Figure 3L and Figure 3M) containing a higher number of myonuclei (Figure 3N and Figure 3O) compared to control muscles. Interestingly, quantification of sub-laminar MuSCs did not show any difference between both genotypes at 62 d.p.i., suggesting that *Rspo1* does not control MuSC self-renewal (data not shown). Taken together, our results suggest that *Rspo1* is required for the proper timing of myogenic progenitor cells differentiation following injury and that *Rspo1* deficiency increase muscle cell fusion *in vivo*.

**Figure 3.**
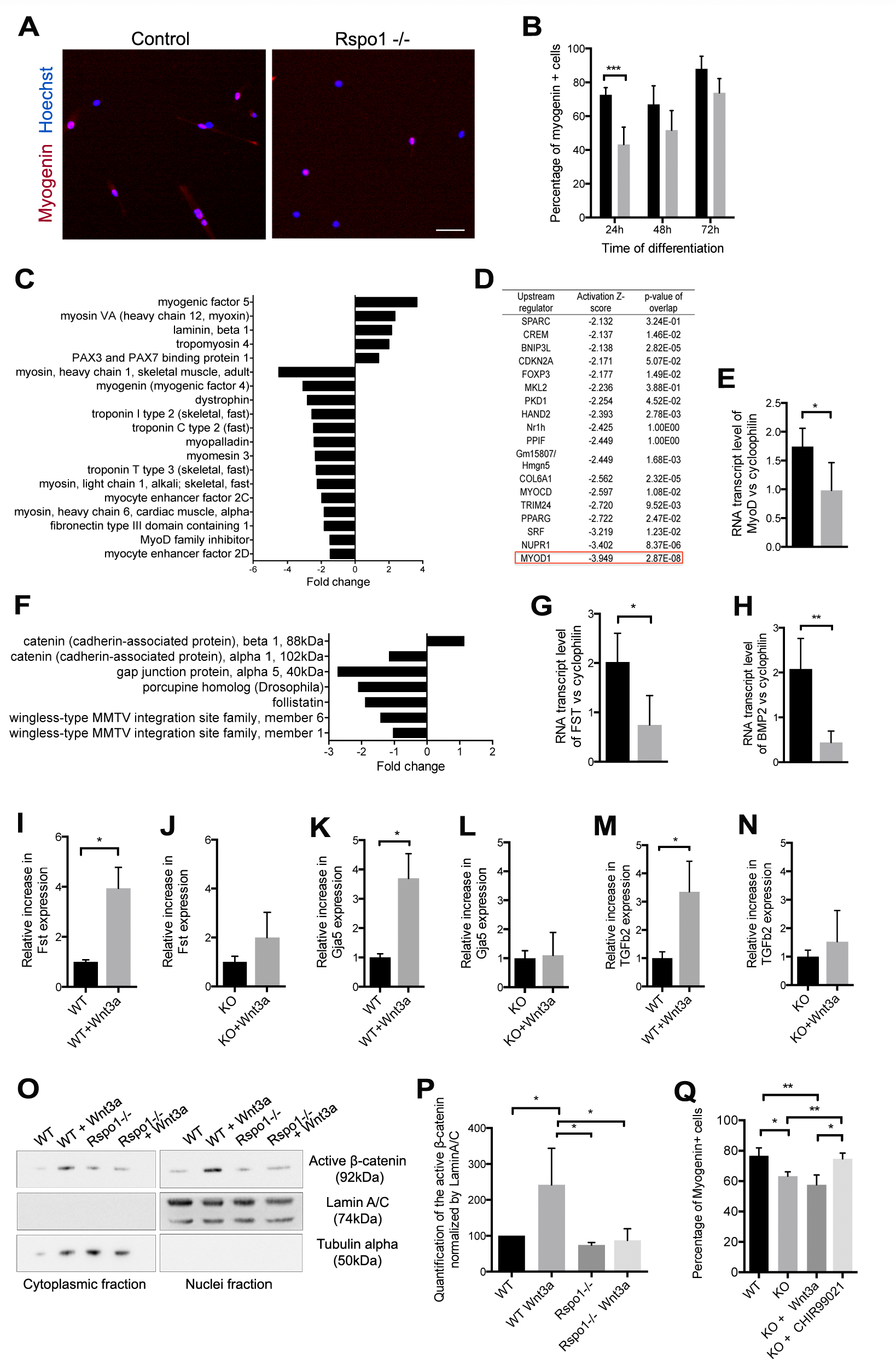
*Rspo1-null* cells have a differentiation defect due to an impaired activation of the Wnt/β-catenin pathway. (A) Immunolocalization of MYOGENIN (red) proteins in control and *Rspo1-null* myocytes. Nuclei are stained with Hoechst (blue). (B) Quantification of differentiated cells. (C) List of genes involved in muscle development and regulated by Rspo1, depending on their fold change in gene expression in *Rspo1-null* versus control myocytes. (D) Upstream Regulators analysis of *Rspo1-null* myocytes transcriptome. (E) qPCR analysis of *MyoD* expression in Rspo1-null and control myoblasts. (F) List of Rspo1-regulated genes involved in canonical Wnt pathway. (G) qPCR analysis of *Follistatin* gene expression in differentiated *Rspo1-null* and control myocytes. (H) qPCR analysis of *Bmp2* gene expression in differentiated *Rspo1-null* and control myocytes. (I,K,M) Increased expression of *Follistatin, Gja5* and *Tgfβ2* following WNT3a treatment in control cells. (J,L,N) Unchanged expression of *Follistatin, Gja5* and *Tgfβ2* following WNT3a treatment in *Rspo1-null* cells. (O) Western blot and (P) quantification of the active β-CATENIN showing a decrease in β-CATENIN protein levels in nuclei of *Rspo1-null* primary myoblasts treated with WNT3a recombinant protein. (Q) Quantification of the number of differentiated cells in control and *Rspo1-null* myocytes, after a WNT3a-or a CHIR99021-treatment, demonstrating that the forced-activation of the Wnt/β-catenin pathway in *Rspo1-null* cells restores a control state of differentiation. Bars: 50 μm in A. *Error bars indicate standard deviation. *: p value <0.05.* **: *p value <0.01.* *** p value <0.001

### *Rspo1* does not influence primary myoblasts proliferation

To further analyze the behavior of muscle cells devoid of *Rspo1*, we cultured primary myoblasts expanded from MACS-sorted MuSCs. We observed that *Rspo1-null* cells do not show an aberrant morphology and expressed the muscle progenitor marker M-cadherin (Figure S2A) (Cornelison and Wold, 1997). While proliferating, control and *Rspo1-null* myoblasts expressed similar levels of *Pax7* transcript expression (Figure S2C) and more than 98% of the cells expressed PAX7 protein (Figure S2B). Immunostainings for the mitosis marker phospho-HistoneH3 (Figure 4D) and the cell cycle marker MKI67 (Figure S2E) did not reveal any differences in the number of cycling cells between *Rspo1-null* and control cells. Likewise, quantification of the proportion of cells engaged in S phase by EdU incorporation assay (Figure S2F) did not show any difference between control and *Rspo1-null* cells in either high serum (Figure S2G) or low serum (Figure S2H) culture conditions. Our results demonstrate that *Rspo1* is not required for myogenic progenitor expansion from MuSCs.

**Figure 4.**
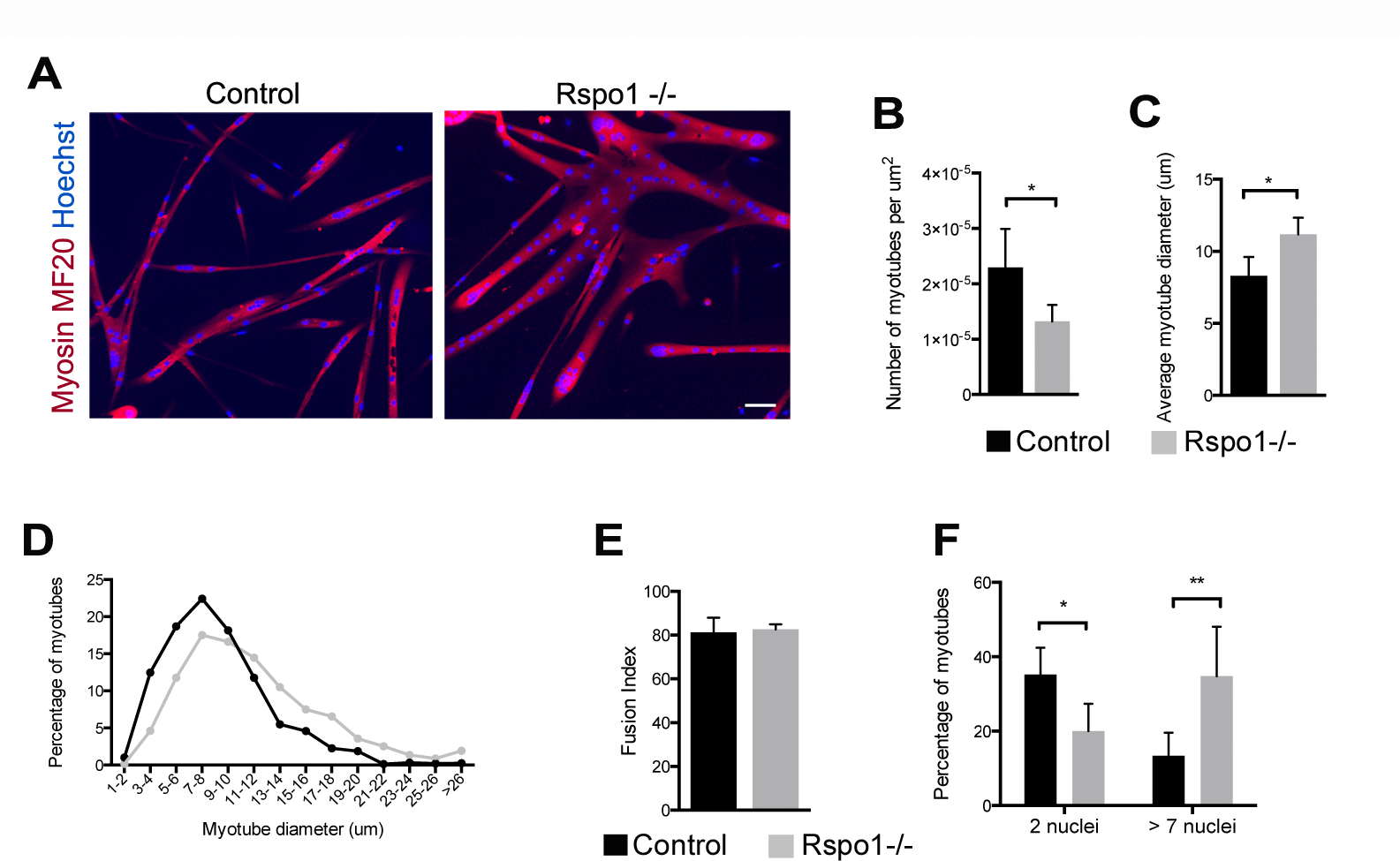
*Rspo1-null* cells show enhanced fusion. (A) Immunolocalization of MYOSIN HEAVY CHAINS (MF20) proteins (red) in control and *Rspo1-null* myotubes after 4 days of differentiation. Nuclei are stained with Hoechst (blue). (B) Quantification of the number of myotubes per surface. (C) Mean diameter of myotubes showing that *Rspo1-null* myotubes are 30% bigger than control ones. (D) Distribution of the percentage of myotubes depending on their diameter size. (E) Quantification of the number of fused cells normalized by the total number of cells. (F) Quantification of the number of nuclei per myotubes showing a decrease proportion of *Rspo1-null* myotubes with 2 nuclei and an increase number of *Rspo1-null* myotubes with a higher number of nuclei. Bars: 50 μm. *Error bars indicate standard deviation. *: p value <0.05.* **: *p value <0.01.* *** p value <0.001

### *Rspo1* positively controls muscle cell differentiation through the Wnt/p-catenin pathway

We further induced *Rspo1-null* and control primary myoblasts to differentiate *in vitro* and quantified the number of myocytes expressing Myogenin at different time points following serum removal (Figure 5A). Similar to our *in vivo* observations, *Rspo1-null* myocytes showed a significant differentiation delay compared to control myocytes, and this delay was compensated after 2 days in differentiation medium (Figure 5B). To understand the molecular mechanisms regulated by Rspo1 in muscle cells, we performed Affymetrix microarrays on *Rspo1-null* and control myocytes. Analysis of the transcriptome data identified the down-regulation of a large number of genes implicated in skeletal myogenesis in *Rspo1-null* cells compared to control cells (Figure 5C). By analyzing our microarray data with the Upstream Regulators tool of the Ingenuity Pathway Analysis (IPA) software, we identified the myogenic determination factor MYOD1, as the most inhibited regulator in *Rspo1-null* cells (Figure 5D). We then validated the down-regulation of *MyoD* transcripts by qRT-PCR (Figure 5E) thus explaining the delay in differentiation of *Rspo1-null* muscle progenitor cells observed *in vitro* and *in vivo.*

**Figure 5.**
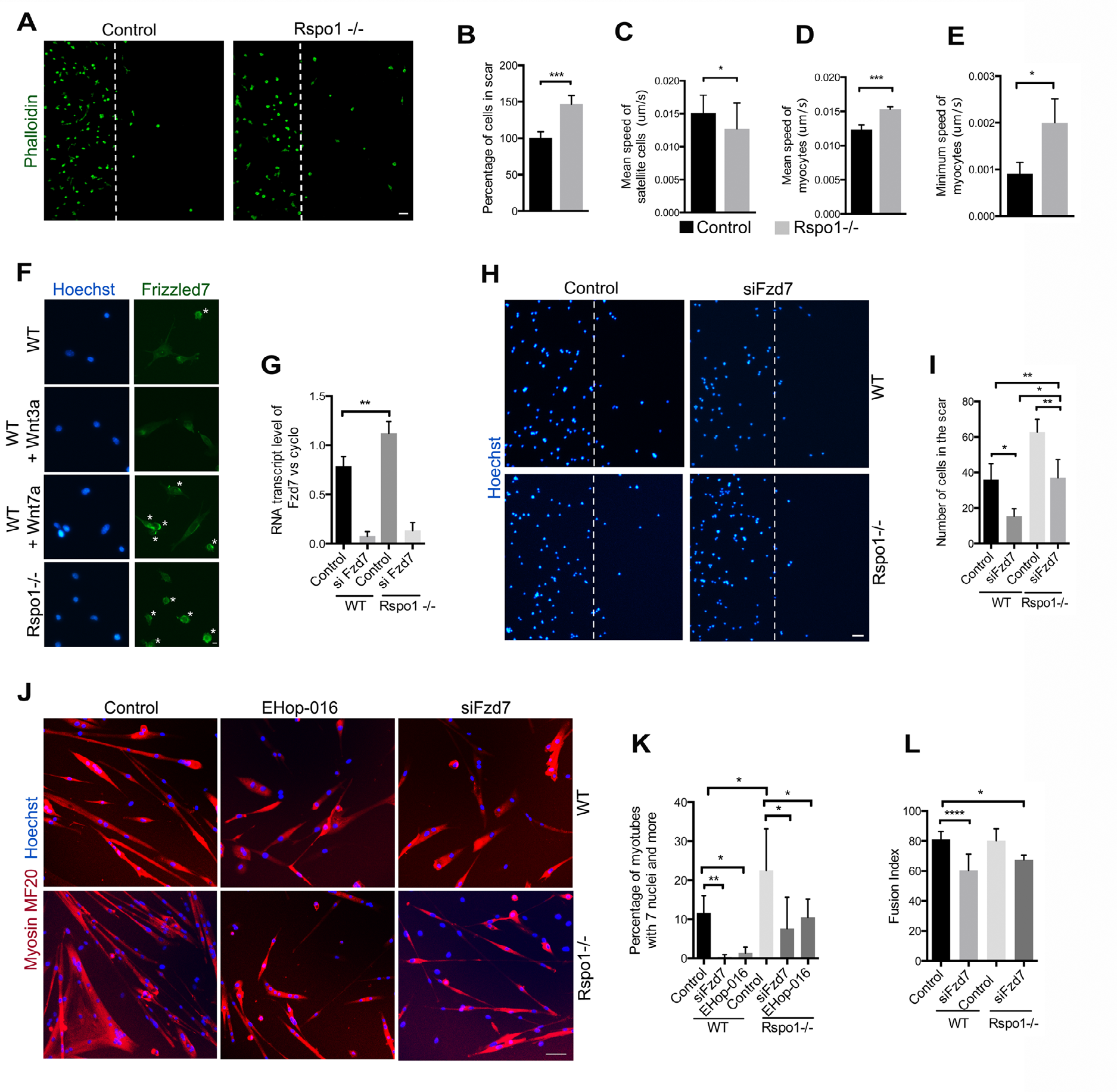
*Rspo1* depletion potentiates Wnt7a/Fzd7/Rac1 signaling. (A) Phalloidin (green) staining of control and *Rspo1-null* myocytes, 24h after scratching confluent-state cells. White bar show the limits of the scratch. (B) Quantification of the number of cells in the scar, 24h after the scratch, showing increases migration of *Rspo1-null* mycytes. (C) Quantification of the mean speed of MuSCs on single myofiber. (D) Quantification of the mean speed and of (E) the minimum speed of myocytes, showing *Rspo1-null* cells have increased velocity. (F) Immunostaining of FRIZZLED7 (green) in *Rspo1-null* and in control myocytes treated by WNT3A and WNT7A recombinant proteins. Nuclei are stained with Hoechst (blue). Asterix represent FRIZZLED proteins located at the cell membrane. (G) qPCR analysis of *Fzd7* expression in myocytes confirming siRNA efficiency and that Rspo1 depletion results in increased *Fzd7* expression. (H) Hoechst stainings of control and *Rspo1-null* myocytes, 24h after scratching confluent-state cells, upon siFzd7 treatment. White bar show the limits of the scratch. (I) Quantification of the number of migrating cells showing a decreased migration upon *Fzd7* silencing. (J) Immunostaining of MYOSIN HEAVY CHAINS (MF20) (red) in control and *Rspo1-null* myotubes treated with siFzd7 or EHop16. Nuclei are stained with Hoechst (blue). (K) Quantification of the number of myotubes with more than 7 nuclei. (L) Fusion index of control and *Rspo1-null* myotubes upon siFzd7 treatment. Bars: 50 “m in A,H,J; 20 “m in F. *: p value <0.05. ** p value <0.01. *** p value <0.001. ****: p value <0.0001. Error bars correspond to standard deviation

The IPA tool further highlighted a down-regulation in the expression of a set of genes implicated in canonical Wnt pathway in *Rspo1-null* cells (Figure 5F). We validated that the expression levels of the μ-catenin target genes *Follistatin* (Fst) (Figure 5G) and *Bmp2* (Figure 5H) were decreased in *Rspo1-null* myocytes compared to control cells. We then investigated if Wnt/β-catenin target genes expressions could be induced in *Rspo1-null* cells following exogenous ligand stimulation. Following WNT3A protein treatment, control myocytes exhibited elevated expression of *Fst* (Figure 5I), *Gja5* (Figure 5K) and *Tgfβ2* (Figure 5M), three genes that we previously showed to be responsive to β-catenin activation in muscle cells (Rudolf et al., 2016). In contrast, *Rspo1-null* myocytes did not respond to exogenous WNT3A stimulation, as expression levels of *Fst* (Figure 5J), *Gja5* (Figure 5L) and *Tgfβ2* (Figure 5N) remained similar in WNT3A-treated cells and in mock-treated cells. We then quantified the expression of active-β-catenin proteins in nuclear and cytoplasmic fractions by Western Blot (Figure 5O). We observed that while active-β-catenin proteins are elevated and translocated to the nuclei of control cells following Wnt ligand stimulation, they remain unchanged in *Rspo1-null* cells following WNT3A treatment (Figure 5P). Since activation of Wnt/β-catenin signaling is required in muscle progenitor cells to properly differentiate (Rudolf et al., 2016), we hypothesize that a lack of β-catenin activation in *Rspo1-null* myocytes could explain the observed delay in differentiation. To stabilize β-catenin proteins in muscle cells, independently of the extracellular ligands, we used the small molecule CHIR99021 to inhibit GSK3 and subsequently the β-catenin destruction complex. Quantification of the number of differentiated cells in control and in *Rspo1-null* myocytes, after treatment with either WNT3A or CHIR99021, showed that the forced-activation of the Wnt/β-catenin pathway in *Rspo1-null* cells restore a control state of differentiation. Taken together, our data suggest that Rspo1 expression is required for the activation of the canonical Wnt pathway during myogenic differentiation.

### *Rspo1* limits cell fusion

We next investigated muscle cell fusion in *Rspo1-null* and control primary myocytes. After four days in differentiating conditions, control myocytes fused and generated both thin and large myotubes, while *Rspo1-null* cells generated larger syncytia (Figure 4A). As such, the number of myotubes per surface was reduced in *Rspo1-null* cultures, compared to control cells (Figure 4B) and the average diameter of *Rspo1-null* myotubes was higher compared to control myotubes (Figure 4). Graphical distribution of myotubes size further demonstrated that this increase was homogenous and not due to the appearance of a specific sub-population (Figure 4D). Importantly, while the percentage of nuclei that underwent fusion was similar in cultures of both genotypes (Figure 4E), we observed a significantly higher proportion of myotubes containing more than 7 nuclei in *Rspo1-null* cultures compared to the control cultures (Figure 4F). Our results indicate that *Rspo1* negatively regulates muscle cell fusion and that its absence leads to the generation of larger myotubes containing more nuclei.

### *Rspo1* negatively regulates the non-canonical Wnt7a/Fzd7/Rac1 pathway

Our data suggests that Rspo1 has contradicting roles in muscle progenitor cell differentiation and fusion. We thus hypothesized that *Rspo1* depletion could also impact the non-canonical Wnt pathway. It is known that WNT7A promotes the activation of the Wnt/PCP pathway in MuSCs (Le Grand et al., 2009). It was also recently shown that WNT7A stimulate the migration of MuSCs and primary myoblasts through RAC1 activation (Bentzinger et al., 2014). We thus validated, that WNT7A stimulation of wild-type differentiating myocytes results in enhancement of muscle fusion, as shown by increased myotube size (Figure S3A) and higher numbers of myonuclei (Figure S3B) in WNT7A-stimulated myotubes compared to mock-treated cells. As such, up-regulation of non-canonical Wnt7a signaling could explain the increased fusion we observed in *Rspo1-null* muscle cells.

To test this hypothesis, we first quantified the migration of control and *Rspo1-null* myocytes in scratch-wound assays (Figure 5A). We observed that *Rspo1* deficiency increased myocyte migration (Figure 5B). To test the effect of *Rspo1* on MuSC migration, we then used time-lapse imaging on single myofibers (Otto et al., 2011). In this condition, the mean velocity of *Rspo1-null* MuSCs was lower than the mean velocity of control cells (Figure 5C). Interestingly, time-lapse imaging of myocytes migrating in wound assays revealed that *Rspo1* deficiency significantly increased both the mean velocity (Figure 5D) and the minimum speed (Figure 5E) when compared with controls. These results indicate that *Rspo1* inhibits the motility of myogenic progenitor cells, but not MuSCs, during directed migration.

The major receptor of WNT7A in myogenic cells is *Frizzled-7* (*Fzd7*) (Le Grand et al., 2009) and it accumulates in the periphery of moving cells during migration (Bentzinger et al., 2014). We thus performed immunolocalization of FZD7 in primary myocytes, and observed that while few cells show FZD7 accumulation in control and WNT3A-treated conditions, FZD7 staining appeared stronger and polarized in both WNT7A-treated cells and *Rspo1-null* cells (Figure 5F). We also observed an increase in *Fzd7* gene expression in *Rspo1-null* cells compared to control cells (Figure 5G). Both these results suggest an enhancement of non-canonical Wnt7a/Fzd7 signaling in *Rspo1-null* cells.

We next transfected both control and *Rspo1-null* cells with siRNA directed against *Fzd7* in order to block the non-canonical WNT7A pathway (Figure 5G). We then quantified the number of migrating cells in scratch wound assays (Figure 5H) and observed that siFzd7 treatment decreased cell migration in both control and *Rspo1-null* cells. Importantly, *Rspo1-null* cells treated with siFzd7 migrated with a rate similar to control cells, indicating that the increase in cell motility observed in *Rspo1-null* cells is related to an increase in Wnt7a/Fzd7 signaling (Figure 5I). To further investigate whether the improved muscle cell fusion resulting from *Rspo1* deficiency is dependent on up-regulation of Wnt7a/Fzd7/Rac1 signaling, we differentiated control and *Rspo1-null* myocytes, in control and *Fzd7* silencing conditions or in the presence of RAC1 inhibitor EHop-016 (Montalvo-Ortiz et al., 2012) (Figure 5J). Strikingly, treatment of *Rspo1-null* cells with either siFzd7 or EHop-16 restored the proportion of large myotubes containing more than 7 nuclei to a percentage similar to untreated control cells (Figure 5K). Interestingly, siFzd7 treatment resulted in a small reduction in fusion index in both controls and *Rspo1-null* cells (Figure 5L).

Taken together, our results indicate that *Rspo1* negatively regulates muscle cell migration and fusion by dampening non-canonical Wnt7a/Fzd7/Rac1 pathway. Our data suggest that, in MuSCs progeny, *Rspo1* acts as a regulator of both canonical and non-canonical Wnt signaling pathways, and that it integrates antagonistic pathways for fine-tuning of muscle architecture during tissue repair.

## DISCUSSION

In most mammalian tissues, the R-spondin family of secreted proteins positively regulates the canonical Wnt signaling pathway (Carmon et al., 2011) by neutralizing Rnf43 and Znrf3, two transmembrane E3 ligases that remove Wnt receptors from the cell surface (de Lau et al., 2014). Canonical Wnt signaling plays an important role in adult muscle regeneration, through allowing for the timely differentiation of MuSC descendants during tissue repair (Figeac and Zammit, 2015, Rudolf et al., 2016). Here we show that the absence of *Rspo1* expression significantly affects myogenic progenitor cells differentiation both *in vitro* and *in vivo*, and is associated with a defect in canonical Wnt/β-catenin activation. These results are consistent with previous reports demonstrating that RSPO1 is a regulator of tissue-resident stem cells such as the mammary gland (Cai et al., 2014), the skin (Li et al., 2016) or the intestine (de Lau et al., 2011).

Following injury, *Rspo1-null* muscle progenitor cells show a transient delay in the acquisition of the differentiated state. They further overtake this deficiency, probably due to the compensatory effects of other signaling pathways. Strikingly, we observed that *Rspo1-null* muscles can further regenerate efficiently and are composed of bigger myofibers containing higher numbers of myonuclei. This increase in fusion is due to an up-regulation of non-canonical Wnt7a/Fzd7/Rac1 signaling pathway in muscle progenitor cells. As such, *Rspo1-null* muscle cells migrated faster and fused with higher efficiency compared to wild-type cells, and these phenotypes were restored following *Fzd7* silencing or RAC1 inhibition. Importantly, while it has been shown that RSPO3 can activate the non-canonical Wnt pathway during *Xenopus* embryonic development (Ohkawara et al., 2011), the impact of R-spondins on non-canonical Wnt pathways has not been previously demonstrated in adult mammalian cells. To date, a dampening function of RSPO1 on the non-canonical Wnt pathway has not been studied, regardless of the tissue or animal species. Further work could then study of the role of R-spondins proteins in controlling non-canonical Wnt pathways in other adult stem cells.

Activation of the Wnt7a/Fzd7 pathway in muscle satellite stem cells induces symmetrical their symmetrical expansion, by activating the Planar Cell Polarity pathway (Le Grand et al., 2009). As such, Wnt7a overexpression enhance muscle regeneration and increase both satellite cell numbers and the proportion of satellite stem cells (Le Grand et al., 2009, von Maltzahn et al., 2012). In contrast, while we observed that *Rspo1* deficiency resulted in over-activation of Wnt7a/Fzd7/Rac1 pathway in differentiating myocytes, we did not quantify any differences in the numbers of PAX7-expressing myogenic progenitors during muscle regeneration or in the MuSCs pool after regeneration is finished. Further to that point, while *Rspo1-null* myocytes are faster than control cells, *Rspo1-null* MuSC on single fiber are not. Our data suggests that RSPO1 does not influence the non-canonical Wnt pathway in MuSC. We hypothesize that this stage-specific requirement for RSPO1 action is related to the fact that canonical Wnt/β-catenin signaling is inactive in MuSC, but activated in differentiating myogenic progenitor cells (Rudolf et al., 2016). We propose that *Rspo1* controls the antagonistic balance between canonical and non-canonical Wnt pathways specifically in differentiating muscle progenitor cells.

Both canonical and non-canonical Wnt signalings induce distinct cellular and molecular processes but share several core components. Thus, the exclusive activation of a Wnt pathway is possible by the selective interaction between specific ligands and receptors. More precisely, it has been shown that WNT5a can inhibit or activate β-catenin signaling depending on the presence of specific Frizzled receptors at the membrane (Mikels and Nusse, 2006). It is also well documented that canonical and non-canonical Wnt ligands can compete for the activation of their selective pathways. In Xenopus embryos, ectopic non-canonical WNT5a can antagonize canonical WNT8 and β-catenin activation (Larabell et al., 1997). In mammals, non-canonical WNT5a has been shown to inhibit WNT3a-mediated canonical Wnt signaling in hematopoietic stem cells (Nemeth et al., 2007). The increase in non-canonical Wnt signaling observed in *Rspo1-null* muscle cells could then be related to a change in the composition of surface receptors. We propose that RSPO1 has a role in maintaining the “canonical” Frizzled receptors at the surface of muscle cells, and that its absence can lead to a preferential increase in the availability of non-canonical Frizzled receptors. The increase in Fzd7 expression in *Rspo1-null* cells may start answering this question, but further work will be dedicated in exploring the expression pattern of the different Frizzled expressed at the surface of MuSCs and their descendants, their dynamics during myogenic commitment and myotube formation.

Our data show that *Rspo1-null* muscles can regenerate efficiently and show increase muscle progenitor cell fusion leading to larger syncytia containing more myonuclei compared to control muscles. As such, blocking RSPO1 function in vivo in myodegenerative diseases or in conditions of muscle atrophy due to ageing or cancer-driven cachexia may represent an interesting strategy for enhancing intrinsic muscle repair. In contrast, the regeneration of bigger myofibers can also be described as a pathological context since it can perturb muscle homeostasis and function. An example is the hyper-muscular *Myostatin-null* mice, in which lack of MYOSTATIN promotes growth of skeletal muscle, but compromises force production (Amthor et al., 2007). Precisely, muscles from *Myostatin-null* mice, although larger and stronger, fatigue extremely rapidly (Mouisel et al., 2014). As such, delivery of RSPO1 protein could serve as a therapeutic treatment in muscle pathologies that show aberrant or elevated muscle progenitor fusion such as Duchenne Muscle Dystrophy.

## METHODS Mice

### Mice

Experimental animal protocols were performed in accordance with the guidelines of the French Veterinary Department and approved by the University Paris-Descartes Ethical Committee for Animal Experimentation. The R-spondin1 knock-out mice were a gift from M-C. Chaboissier (Institut de Biologie Valrose, Nice, France). All experiments were performed in 6-to 9-weeks-old mice. Our animals are on C57B6N/SV129 genetic background. Mice genotyping has been described previously (Chassot et al., 2008).

### Cardiotoxin injury

Mice were anaesthetized by intraperitoneal injection of Ketamin at 0,1mg per gram body weight and Xylazin at 0,01mg per gram body weight diluted in saline solution. After having cleaned the mouse hind legs with alcohol, tibialis anterior muscles were injected with 50μl of cardiotoxin solution (Latoxan, 12μM in saline) using an insulin needle.

### Muscle histology and immunohistochemistry

For cryosections, TA were submerged in Tissue-Tek O.C.T. compound from Cell Path. Beforehand, isopentane was cooled in liquid nitrogen. Muscles were then frozen on the cooled isopentane and cut in 10 μm section with a Leica cryostat. Frozen muscle were stored at −80°C. Sections were fixed with 4% PFA in PBS during 20 min and permeabilized with cooled-methanol. Antigen retrieval was performed with the Antigen Unmasking Solution (Vector) at 95°C for 10 min. Sections were then blocked with 4% BSA, 5% Goat serum in PBS during 3h and incubated with primary antibody overnight at 4°C. Alexa Fluor secondary antibody was incubated on sections 1h at room temperature and nuclei were stained with Hoechst. Slides were mounted in fluorescent mounting medium from Dako. Primary antibodies used were Laminin (Santa Cruz), Pax7 (Santa Cruz), Myogenin (Santa Cruz) and Dystrophin (Leica). For Hematoxilin/Eosin staining, sections were incubated in Hematoxilin bath for 4 min and with Eosin for 2 min and thereafter dehydrated in 70%, 90% and 100% ethanol and fixed in Xylene bath for 2 min.

### Muscle stem cells isolation and primary myoblasts culture

Skeletal muscles of mice were dissected (Quadriceps, TA, EDL, Gastrocnemius, Soleus, Gluteus) and transferred to a sterile Petri dish on ice. Muscles were incubated in 1,5U/mL of CollagenaseB - 2,4U/mL of DispaseII - 2M of CaCl_2_ solution for 45min at 37°C with periodic mechanical digestion. Fetal Bovine Serum was added to stop the digestion. After centrifugation, pellet was resuspended in growth medium consisting of Ham’s F10 (Life Technologies) with 20% FBS (Eurobio), 1% Pen/Strep (Life Technologies), 2.5 ng/μl basicFGF (R&D Systems). Satellite cells were then purified using MACS cell separation system, according to the manufacturer’s protocol. Satellite cells are then let to proliferate and give rise to primary myoblasts after two passages. If needed, primary myoblasts were differentiated using a differentiation medium composed of DMEM (Gibco) with 2% Horse Serum (Gibco), 1% Pen/Strep (Gibco). Cells were treated with 50ng/mL of recombinant WNT3A and/or WNT3A proteins (R&D Systems) for a minimum of 24h, or with the RAC1 inhibitor eHop-016 (Sigma) at 1,5μM during 8h. Primary myoblasts overexpressing full-length Pax7 (cloned into the pBrit plasmid vector) were generated by retroviral infection and cultured as previously described (McKinnell et al., 2008).

### Chromatin immunoprecipitation

Primary myoblasts were cross-linked using 1% formaldehyde in PBS 1X for 10 minutes. Glycine was then added to a final concentration of 0,125M for 5 minutes, followed by centrifugation. Cell pellet was washed in PBS 1X, and resuspended in ChIP lysis buffer (50mM Tris-HCl pH 8.0, 10mM EDTA, 0,5% SDS). Chromatin was fragmented using Covaris sonicator. Before ChIP, equal volume of ChIP assay buffer (20mM Tris-HCl pH 8.0, 1,2mM EDTA, 1,1% Triton X-100, 200mM NaCl) was added to adapt the concentration of salts and detergents. Immunoprecipitation was performed using 1mg of chromatin and 30μl of anti-Flag M2 affinity gel (Sigma) for 3h at 4 °C. Antibody-protein-DNA complexes were collected, washed and eluted, and cross links were reversed according to manufacturer's instructions. DNA was purified by phenol/chloroform purification using linear acrylamide (Ambion) and GlycoBlue (Ambion) as carriers. ChIP enrichment was analyzed by quantitative PCR using Mx3000P (Stratagene).

### Immunostaining and EdU incorporation on primary myoblasts

Primary myoblasts were stained for 8min with 4% PFA in PBS then incubated with 0,2% Triton in PBS for 20min at room temperature. Cells were incubated with primary antibodies during 1h at room temperature, followed by several PBS washes and a 1h-incubation with the secondary antibodies. Nuclei were stained with Hoechst. Primary antibodies used were R-spondin1 (Sigma), Phospho-Histone H3 (Cell Signaling), Pax7 (Santa Cruz), Myogenin (Santa Cruz), M-cadherin (BD Biosciences), Myosin Heavy Chains (R&D Systems) and MKI67 (Santa Cruz). EdU incorporation on primary myoblasts was performed with Click-iT EdU Imaging Kits from Invitrogen, according to the manufacturer’s protocol. Cells were incubated with EdU during 1h at 37°C.

### Scratch assay

Cells were treated with 10μg/mL of Mitomycin-C (Sigma) for 8h before scratching the monolayer of cells in a straight line. Plates were washed few times with PBS and primary myoblasts were incubated in growth medium for 24h at 37°C. If needed, WNT7A recombinant proteins (100ng/mL from R&D Systems) were added in the culture medium. Cells were then fixed with 4% PFA - PBS and nuclei were stained with Hoechst. The numbers of nuclei in the scar were counted manually.

### RNA interference

Primary cells were seeded in collagen-coated plate in growth medium without Pen/Strep. SiRNA transfection was performed using Lipofectamin 2000 and OptiMEM according to the manufacturer’s protocol. Both siFzd7 and siControl (Ambion) were used at a 50μM concentration.

### Live imaging

Cell imaging was performed on a microscope (Zeiss Axio Observer.Z1) with x10 objective under 5% CO_2_ at 37°C in differentiation media. Image acquisition was performed with Metamorph software (Molecular Devices) with time points acquired every 8min during 24h. Velocity was calculated with ImageJ software. The minimum velocity is defined as the lowest velocity obtained by cells during the tracking period.

### Quantitative real-time PCR (qPCR)

RNA was isolated either by RNeasy Mini Plus Kit (Qiagen) or by TRIzol Reagent followed by a DNAse treatment (Life Technologies). The reverse transcription was performed with 100ng of RNA using the High-Capacity cDNA Reverse Transcription kit (Applied Biosystems). Transcript levels were determined by LightCycler 480 Real-Time PCR System from Roche, using SYBR green I Master from the same company. The melting curves were checked for each experiment and the primers efficiency was calculated by serial sample dilution. Targeted gene expressions were normalized by Cyclophilin reference gene. Sequences of primers used are given in supplementary materials.

### Microarray and bioinformatics

The RNA from primary myoblasts were isolated using TRIzol Reagent from Life Technologies according to the manufacturer's protocol. The purity and the quality of the RNA were performed by Bioanalyzer 2100, using Agilent RNA6000 nano chip kit (Agilent Technologies) at the Cochin Institute Genomic facility. RNA were reverse transcribed using the Ovation PicoSL WTA System (Nugen). The product cDNA were then hybridized to GeneChip Mouse Gene 2.1 ST (Affymetrix). The Affymetrix data were normalized using R software (R Development Core Team, 2011) and gene expression levels were compared using one-way ANOVA. In R-spondin1 deficiency myocytes, the genes were selected with p<0,05 significance. The functional analysis of the gene expression levels was interrogated with Ingenuity pathway (Ingenuity Systems, http://www.ingenuity.com).

### Western blot analysis

The nuclear and cytoplasmic protein fractions were obtained with NE-PER Nuclear and Cytoplasmic Extraction reagents kit (Thermo Scientific) according to the manufacturer’s protocol. Protein amounts were quantified by BCA Protein Assay Kit (Pearce). In each condition, 30μg of proteins was prepared with home-made Laemmli. Samples were runned on NuPAGE 4–12% BIS-TRIS gels (Life Technologies) and transferred on nitrocellulose membranes. After blocking in 5% milk and 0.1% Tween-20 in TBS, membranes were incubated with primary antibodies overnight at 4°C. Secondary antibodies were then incubated on the membrane for 1h at room temperature. Signals were detected using SuperSignalWest Pico Chemiluminescent Substrate (Thermo Scientific) with ImageQuant TL LAS 4000 from (GE Healthcare). The primary antibodies used were as follows: LaminA/C (Cell Signaling), active β-catenin (Cell Signaling) and alpha-Tubulin (Sigma).

### Image acquisition and quantitative analysis

Image acquisitions were performed at the Cochin Institute and the ICM Imaging Facility in Paris. Immunofluorescent stainings were analyzed with an Olympus BX63F microscope, Zeiss Axio Observer.Z1 microscope and AZ100 Nikon Macroscope from Cochin Institute, and Zeiss Axiovert 200M microscope and Axio Scan.Z1 microscope from the ICM. We also performed acquisition on EVOS FL Cell Imaging System microscope from Myology Institute (Paris). Fluorochromes used were Alexa Fluor 488 and Alexa Fluor 546 from Life Technologies. NIS-Element (Nikon), Metamorph (Molecular Devices), and Photoshop software (Adobe) were used for image acquisition. Quantifications were performed with ImageJ software.

### Statistical analysis

Analysis were performed using 3 to 5 mice and 3 to 6 biological replicates for cell experiments. Statistical analysis were done using GraphPad software (Prism, CA, USA) with unpaired twotailed Student’s t test, the one-way ANOVA or the two-way ANOVA. P< 0.05 was considered statistically significant. Data were expressed as mean±SD.

## ACKNOWLEDGEMENTS

We thank P. Maire and D. Duprez for critical reading of the manuscript. This work was supported by funding from the INSERM, the CNRS, the Agence National de Recherche (ANR Jeune Chercheur, project RPV13010KKA), E-Rare/ERANET (projet RPV14030KKA) and the Association Fran9aise contre les Myopathies/AFM Telethon. F.L was supported by a fellowship from the DIM Biothérapies. The authors acknowledge B. Durel and P. Bourdoncle of the Cochin Imaging Facility and the members of the PICPS Imaging Facility. We thank members of the Cochin Institute genomic facility (S. Jacques and F. Dumont) for their technical and bioinformatics assistance. The authors declare no competing financial interests.

## Supplemental material

Fig. S1 details skeletal muscle tissue in Rspo1 mutant mice. Fig. S2 shows that Rspo1-null primary myoblast can proliferate efficiently. Fig. S3 presents the increase in muscle cell fusion following Wnt7a stimulation. Table S1 contains the PCR primers sequences.

### Accession codes

The data discussed in this publication have been deposited in NCBI's Gene Expression Omnibus and are accessible through GEO Series accession number GSE84016.

